# Brain dynamics predictive of response to psilocybin for treatment-resistant depression

**DOI:** 10.1101/2022.06.30.497950

**Authors:** Jakub Vohryzek, Joana Cabral, Louis-David Lord, Henrique M Fernandes, Leor Roseman, David J Nutt, Robin L Carhart-Harris, Gustavo Deco, Morten L Kringelbach

## Abstract

Psilocybin therapy for depression has started to show promise, yet the underlying causal mechanisms are not currently known. Here we leveraged the differential outcome in responders and non-responders to psilocybin (10mg and 25mg, 7 days apart) therapy for depression - to gain new insights into regions and networks implicated in the restoration of healthy brain dynamics. We used whole-brain modelling to fit the spatiotemporal brain dynamics at rest in both responders and non-responders before treatment. Dynamic sensitivity analysis of systematic perturbation of these models enabled us to identify specific brain regions implicated in a transition from a depressive brain state to a heathy one. Binarizing the sample into treatment responders (>50% reduction in depressive symptoms) versus non-responders enabled us to identify a subset of regions implicated in this change. Interestingly, these regions correlate with in vivo density maps of serotonin receptors 5-HT_2A_ and 5-HT_1A_, which psilocin, the active metabolite of psilocybin, has an appreciable affinity for, and where it acts as a full-to-partial agonist. Serotonergic transmission has long been associated with depression and our findings provide causal mechanistic evidence for the role of brain regions in the recovery from depression via psilocybin.

## Introduction

Behavioral differences between healthy and depressed individuals can sometimes be conspicuous but identifying causal contributions from brain dynamics is more challenging. Discrete global brain states, such as those that pertain to sleep, healthy waking consciousness and the psychedelic state, have their own characteristic spatio-temporal dynamics, involving large-scale spatial communities temporally evolving in transient arrangements (Sadaghiani *et al*., 2015; Vidaurre *et al*., 2016; Deco *et al*., 2019; Kringelbach and Deco, 2020). With recent advancements in non-invasive neuroimaging techniques, it has become possible to describe complex spatio-temporal dynamics in terms of their spatial and temporal information. Still, one of the challenges for systems neuroscience is to understand what the most appropriate description of such dynamics is and how transition between one state to another is made possible.

A common method for characterizing global brain function, involves assessing how activity is temporally correlated across spatially separate brain areas over an entire recording period, defining static and state-specific ‘functional connectomes’ (Bullmore and Sporns, 2009; Amico *et al*., 2017; Gutiérrez-Gómez *et al*., 2020). However, the last decade has brought clear evidence that finer-grained, more dynamic analysis of brain states, can deepen our understanding of their properties and relationship to behavioural states (Hutchison *et al*., 2013; Allen *et al*., 2014; Calhoun *et al*., 2014). There is a growing taxonomy of approaches to characterize the dynamics of functional interactions (Preti, Bolton and Ville, 2016; Bolton *et al*., 2020; Kringelbach *et al*., 2020), from data-driven heuristic clustering methods across time (Hutchison *et al*., 2013; Allen *et al*., 2014; Calhoun *et al*., 2014; Karahanoğlu and Van De Ville, 2015), dynamical systems informed phase-locking approaches (Cabral *et al*., 2017; Lord *et al*., 2019; Vohryzek *et al*., 2020), Hidden Markov Models (Baker *et al*., 2014; Vidaurre, Smith and Woolrich, 2017) to spatio-temporal networks (Griffa *et al*., 2017; Vohryzek *et al*., 2019).

Efforts and methods are advancing for understanding response to neuropharmacological interventions for depression. Understanding the therapeutic actions of interventions promise - not only to shed light onto the mechanistic relationship between various brain states implicated in health and pathology - but also to provide inspiration for the development of new, improved interventions. However, there are considerable practical and ethical challenges for answering mechanistic questions in humans, elevating the use of animal models (with sometimes questionable translational validity) or small clinically relevant populations – where mechanistic testing can interfere with therapeutic procedures (Arbabyazd *et al*., 2020; Perl *et al*., 2020). One potential advance in this direction, is the use of whole-brain modeling - as a tool for understanding pathological changes in neuropsychiatric disorders, and, potentially, for clinical diagnosis and prediction (Kringelbach and Deco, 2020). We are mindful, however, that the predictive power of any model depends on how well it can describe and predict experimental data to which it is fitted (Cabral *et al*., 2017).

The present paper focuses on whole-brain network models where region specific stimulation or excitation can be tested in silico, and used to describe and predict empirical-informed target states (Deco *et al*., 2019) – such the global brain state found in people with intractable depression. These models link regional dynamics with the neuroanatomical structure of the brain to describe the spatio-temporal activity of functional data (Deco and Jirsa, 2012). This approach bypasses the ethical constrains of human or non-human animal experimental settings, enabling many types of stimulation to be tested, in order to evaluate the role of regions and their excitation on transit between states – with relevance to empirical phenomena of interest. The validity of this strategy has previously been demonstrated in the context of sleep and awake states (Deco *et al*., 2019).

Here, we build on this notion of dynamic sensitivity analysis to gain insight into response to psilocybin therapy for treatment resistant depression. We define brain states in terms of spatial subdivisions and their probability of occurrence across time, characterised as Probabilistic Metastable Substates (PMS). These recurrent metastable substates can be characterized by their probability of occurrence. Beyond the quantitative description of brain states, we wish to understand which brain regions play a prominent role in the recovery from depression after treatment with psilocybin (Vohryzek *et al*., 2022).

Using data from a trial of psilocybin-therapy for treatment-resistant depression, the sample was binarized into ‘responders’ and ‘non-responders’ to psilocybin therapy. Empirical fMRI data was collected before and one-day after the second of two psilocybin-therapy dosing sessions. Using parameters from the empirical data, modeled brain states - and stimulation parameters therein, could then be used to predict treatment response, defined as a >50% reduction in symptom severity from baseline - determined at a key 5-week post-treatment endpoint (Carhart-Harris *et al*., 2016).

Psychedelic medicine has shown a promising avenue for treating depression (Daws *et al*., 2022). For depression treatment, one current hypothesis is that: via a psychedelic drug x psychological intervention combination, there is an increase in global brain flexibility, translating into a window of opportunity for breaking free of negative cognitive biases and associated ruminations (Carhart-Harris and Goodwin, 2017). Indeed, the current research on the acute effects of psychedelic drugs suggests an increase in the repertoire of brain activity substates (Tagliazucchi *et al*., 2014; Atasoy *et al*., 2017; Parker Singleton *et al*., 2021). From a neuropharmacological perspective, psilocybin – an active compound in magic mushrooms – binds with high affinity to the serotonergic 5HT_2a_ receptors but other serotonergic receptors are also implicated (Calvey and Howells, 2018; Carhart-Harris, 2019). Psilocybin acts as an agonist resulting in higher neuronal excitability, modulating the excitatory-inhibitory balance (in favour of excitation) in the cortical brain regions with more 5HT_2a_ receptors (Nutt, King and Nichols, 2013). Recently, a whole-brain computational study focusing on the human brain action of lysergic acid diethylamide (LSD) – which has a similar pharmacology to psilocybin/psilocin – demonstrated, for the first time, the causal impact of 5HT_2a_ agonism-induced excitation on global brain dynamics (Deco, Cruzat, *et al*., 2018).

Here, in empirical fMRI data, we identified recurrent brain substates in terms of the PMS space across all the subjects in the pre- and post-treatment conditions. Furthermore, we use a computational whole-brain model – where each brain area is represented by a Hopf-bifurcation model (Deco, Kringelbach, *et al*., 2017) - to simulate the brain network dynamics in patients before the treatment. Through dynamic sensitivity analysis, we were able to identify brain regions responsible for treatment response at a key 5-week endpoint (Deco, Cabral, *et al*., 2018; Deco *et al*., 2019). A priori, we hypothesised that regions permitting transition to a healthy brain state (as predicted by the 5-week endpoint) would relate to the distribution of the 5HT_2a_ and 5HT_1a_ receptors in the human brain, as determined by prior in vivo positron emission tomography (PET) mapping (Beliveau *et al*., 2017).

## Results

In summary, a quantitative characterization of the spatio-temporal dynamics recorded with fMRI was obtained using leading eigenvector dynamics analysis (LEiDA), resulting in the definition of Probabilistic Metastable Substates (PMS), whose probability of occurrence was compared across conditions (i.e., within-subjects design – therefore, before versus after treatment). We then constructed two whole-brain models representative of the pre-treatment brains to psilocybin therapy. This was done by fitting their PMS descriptions to those obtained from the experimental data. Finally, a dynamic sensitivity analysis was implemented to both responder and non-responder pre-treatment models to identify the brain regions that permit a transition to the healthy PMS (described by responders’ (as predicted by the 5-week endpoint) one-day post treatment brains).

As described in the methods section, we computed the PMS pre- and post-treatment with psilocybin (where ‘post’ = 1 day post psilocybin dosing session two), for both responders and non-responders (determined 5 weeks hence). Here, we focused on a three-substate solution – the lowest k-level with statistically significant differences between the two groups as well as optimal quality measures across clustering solutions (SI Figure 2). When contrasting responders versus non-responders, the occurrence of substate 3 was significantly different pre-versus post-treatment (p = 0.0258, signed rank-sum test), as well as in the post-treatment data alone (p = 0.0141, rank-sum test; Figure 2, A). Furthermore, we also computed the Global Brain Connectivity (GBC), metastability and Functional Connectivity Dynamics (FCD) measures (see SI Figure 2). These results clearly indicated the necessity of considering both spatial and temporal dimensions to differentiate between conditions as GBC, synchrony and metastability show non-significant results. Conversely, the FCD measure showed significant differences in the temporal similarities of spatial patterns between pre- and post-treatment responders (p = 0.0163, signed rank-sum permutation test), and pre- and post-treatment non-responders with post-treatment responders respectively (p = 0.0183 and p = 0.0273, rank-sum permutation test), further supporting the use of spatio-temporal measures to capture the alterations in whole-brain dynamics across conditions.

**Figure 1.**
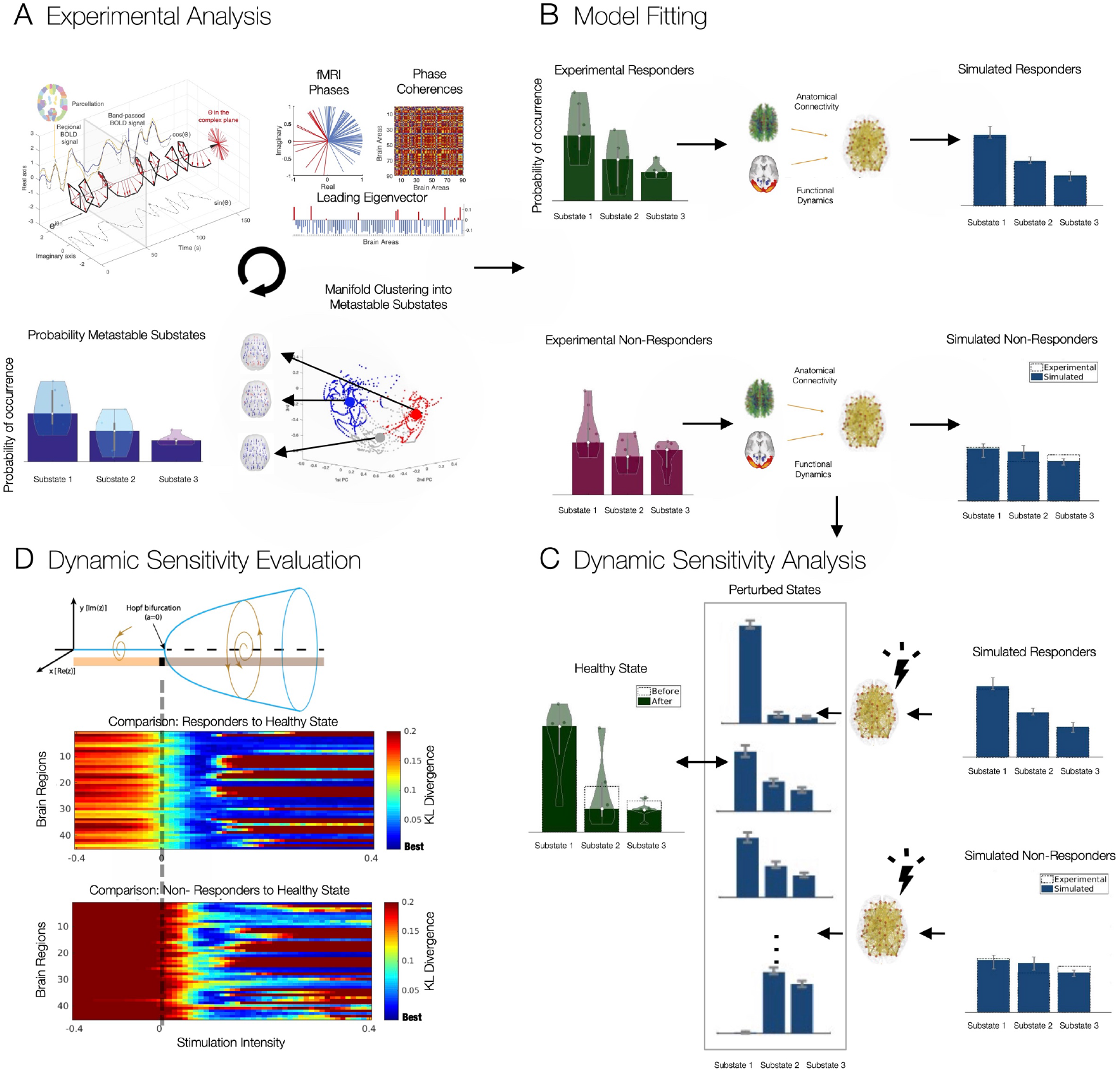
Study Overview. **A) Experimental Analysis.** Probabilistic Metastable Substates were computed for each condition using leading eigenvector dynamics analysis (LEiDA). Regional fMRI timeseries were first converted to analytical signal, followed by computation of the leading eigenvector of the phase coherence matrix at every timepoint. An unsupervised k-means algorithm was deployed to cluster the eigenvectors into a three substate solution. The PMS is defined as the probability distribution of substates, obtained for each individual scan and averaged within each condition. **B) Model Fitting**. Whole-brain model parameters were optimised to fit the PMS before treatment separately for responders and non-responders. **C) Dynamic Sensitivity Analysis**. In silico bilateral perturbations were performed to find the optimal protocol to transition to the PMS characteristic of a healthy brain state (described by responders’ (as predicted by the 5-week QIDS endpoint) one-day post-treatment brains). **D) Dynamic Sensitivity Evaluation**. Perturbations are applied separately in each pair of bilateral brain regions by varying the intensity of oscillations as defined by the bifurcation parameter a.

**Figure 2.**
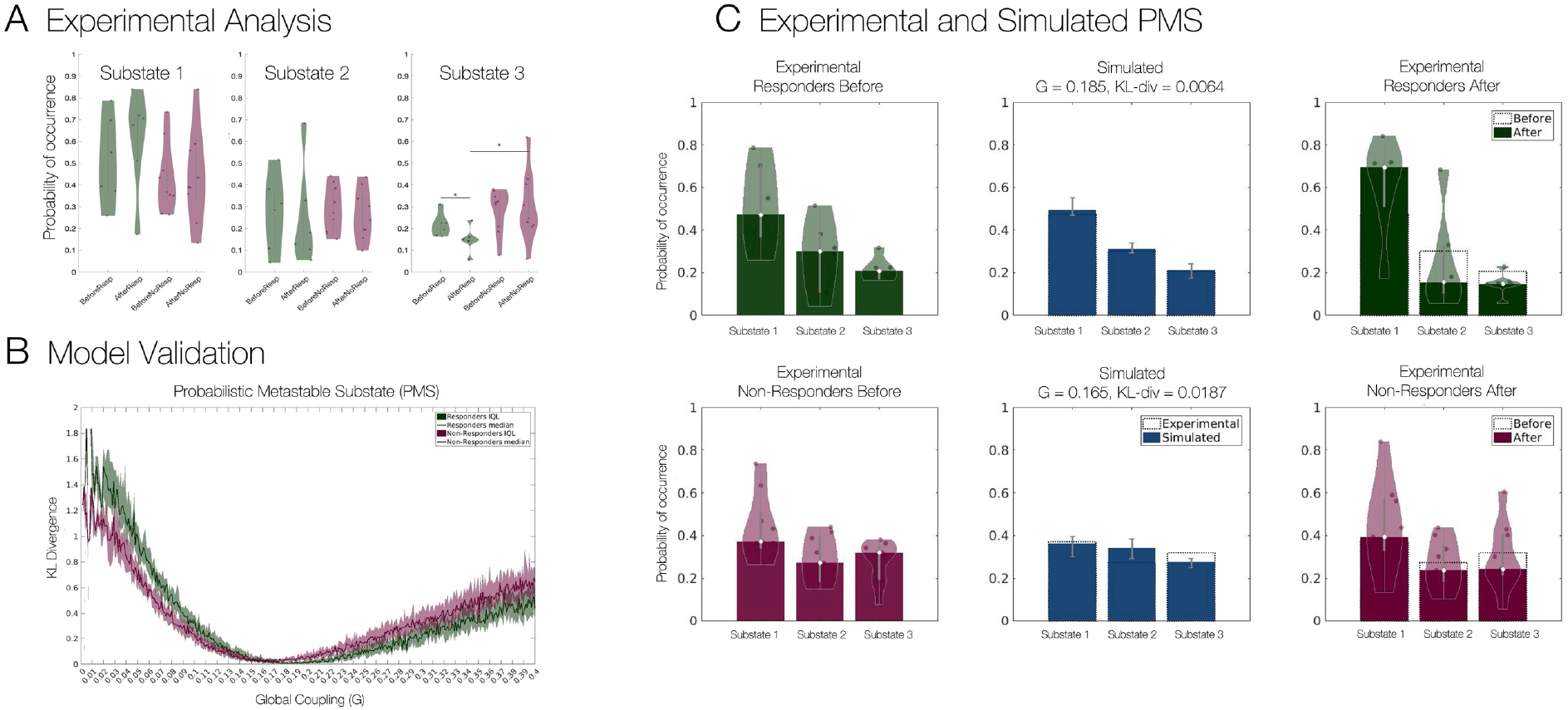
**A) Experimental Analysis.** Probability of occurrence (or Fractional Occupancy) of each metastable substate detected with LEiDA for the three-substate clustering solution. Significant differences were observed in the substate 3 between responders before and after treatment (p = 0.0258, signed rank-sum permutation test), responders and non-responders after treatment (p = 0.0141, rank-sum permutation test) and no significant differences were found between responders and non-responders before treatment. **B) Model Fitting** of the responder and non-responder models as a function of the global coupling parameter G, with optimal fits at G = 0.185 (Kullback-Liebler divergence = 0.0064) and G = 0.165 (KL-divergence = 0.0187) respectively. **C) Experimental and Simulated PMS**. Experimental PMS for responders and non-responders before treatment (left), their simulated counterparts at optimal G (middle), and experimental PMS for responders and non-responders after treatment (right).

To obtain whole-brain computational models representative of the two groups of patients (responders and non-responders before treatment), we first defined a generalized brain network model, where each of 90 cortical and subcortical brain regions (defined using automated anatomical labelling (Tzourio-Mazoyer *et al*., 2002)) was described by a Stuart-Landau oscillator (see methods), and regions were coupled according to realistic structural connectivity obtained from diffusion MRI.

To adjust the model to each group of patients, first the intrinsic frequency of each brain region was set to the peak frequency in fMRI signals averaged across patients in the same group (see SI Figure 3). Subsequently, the global coupling parameter, *G*, was tuned to optimize each model to its appropriate working point. This was achieved by minimizing the divergence between the experimental and simulated PMS spaces -see Figure 2 B. In SI Figure 4, we report optimisation curves for other observables such as the static FC, metastability and FCD. For the responders and non-responders before treatment, we found *G* = 0.185 (KL divergence = 0.0064) and *G* = 0.165 (KL divergence = 0.0187) respectively to minimise the difference. Figure 2 C shows on the left, the experimental results for both groups before treatment; in the middle, the optimal simulated fits for both groups, and on the right the experimental results after treatment (with the results of responders after treatment serving as the target PMS for rebalancing).

Subsequently, we considered a dynamic sensitivity analysis to determine the optimal perturbation strategies to rebalance the PMS distribution to the healthy state (as defined by the PMS space of responders after one-day after treatment). Figure 3 illustrates the dynamic sensitivity analysis, whereby the bifurcation parameter *a* is used to change the nodal dynamics in terms of its response to added noise, ranging from a more noise-driven regime (the more **a** is negative) to an oscillatory regime (with larger amplitude the more **a** is positive). We focused on homological nodal perturbation of the whole-brain model, meaning that bilateral regions were perturbed equally, resulting in 45 pairs of regions perturbed at gradually varying values of *a*. Figure 3, Left, shows the dynamic sensitivity analysis of driving a transition to the healthy state for models of both responders and non-responders before treatment. Again, an average of the KL divergence between either the perturbed pre-treatment responders or non-responder models and the healthy PMS space was shown. In the noise-driven regime (*a* < 0), a deterioration of the fit was observed for both groups, while in the oscillatory regime (*a* > 0), an initial improvement across all 45 runs was depicted, before subsequent deterioration away from the optimal fit for both groups. Conversely, when replacing the target healthy state by the depressive state (i.e., by comparing with the average PMS in non-responders after treatment), we found that the KL divergence was minimal without perturbation (i.e., keeping a=0), showing a worse fit for both groups when brain areas became more oscillatory and no effect of the noisy perturbation (Figure 3, Right).

**Figure 3.**
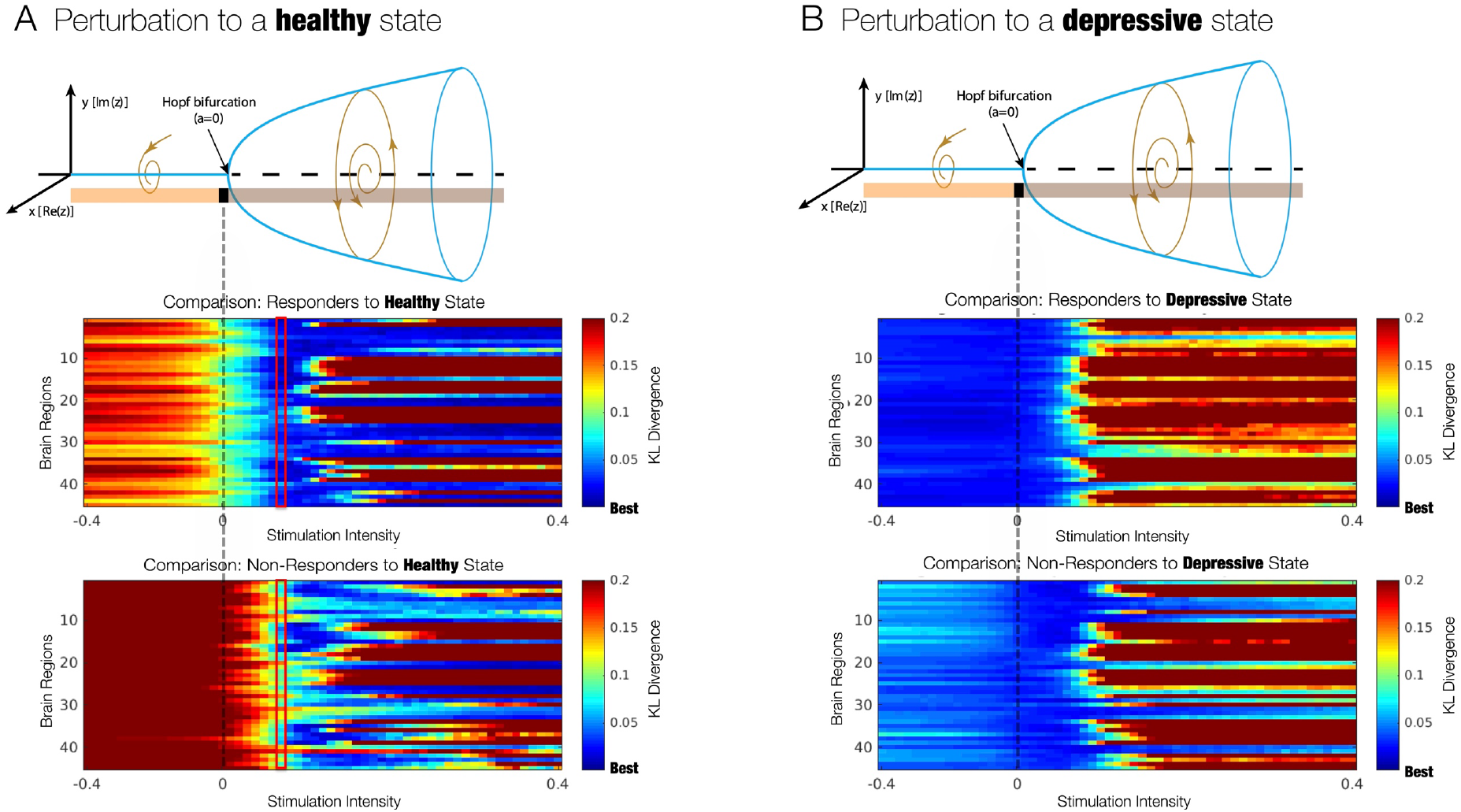
Evaluation of Dynamic Sensitivity Analysis. **A) Perturbation to induce a transition to a healthy state.** Each homological pair of brain regions was perturbed by varying the bifurcation parameter **a**, which modulates the intrinsic oscillatory behavior of the dynamical units. The more **a** is positive, the larger the amplitude of intrinsic oscillations, whereas for negative **a** the units decay to a fixed point equilibrium and the local dynamics is dominated by noise. The performance of the perturbations is evaluated by computing the KL divergence between the simulated PMS and the empirical PMS from patients who recovered after treatment with psilocybin. Optimal intensity of **a** = 0.07 was achieved for the responder group (red rectangles). **B) Perturbation to induce a transition to a depressive state**. A transition to the depressive state showed worse or no effect at varying values of the bifurcation parameter **a**. This is expected since the models were adjusted to patients in the depressive state before treatment.

To evaluate which regions permitted transition to a healthy state, we first defined the optimal perturbation strength as the minimum of the averaged KL divergence (across the 45 runs) of the responder group to the treatment. This stimulation intensity was found at *a* = 0.07. Then, we inspected the difference between the responders and non-responders at that given value of *a* to assess what nodal perturbations were permitting the transition to the healthy state in responders but not in non-responders (Figure 4).

**Figure 4.**
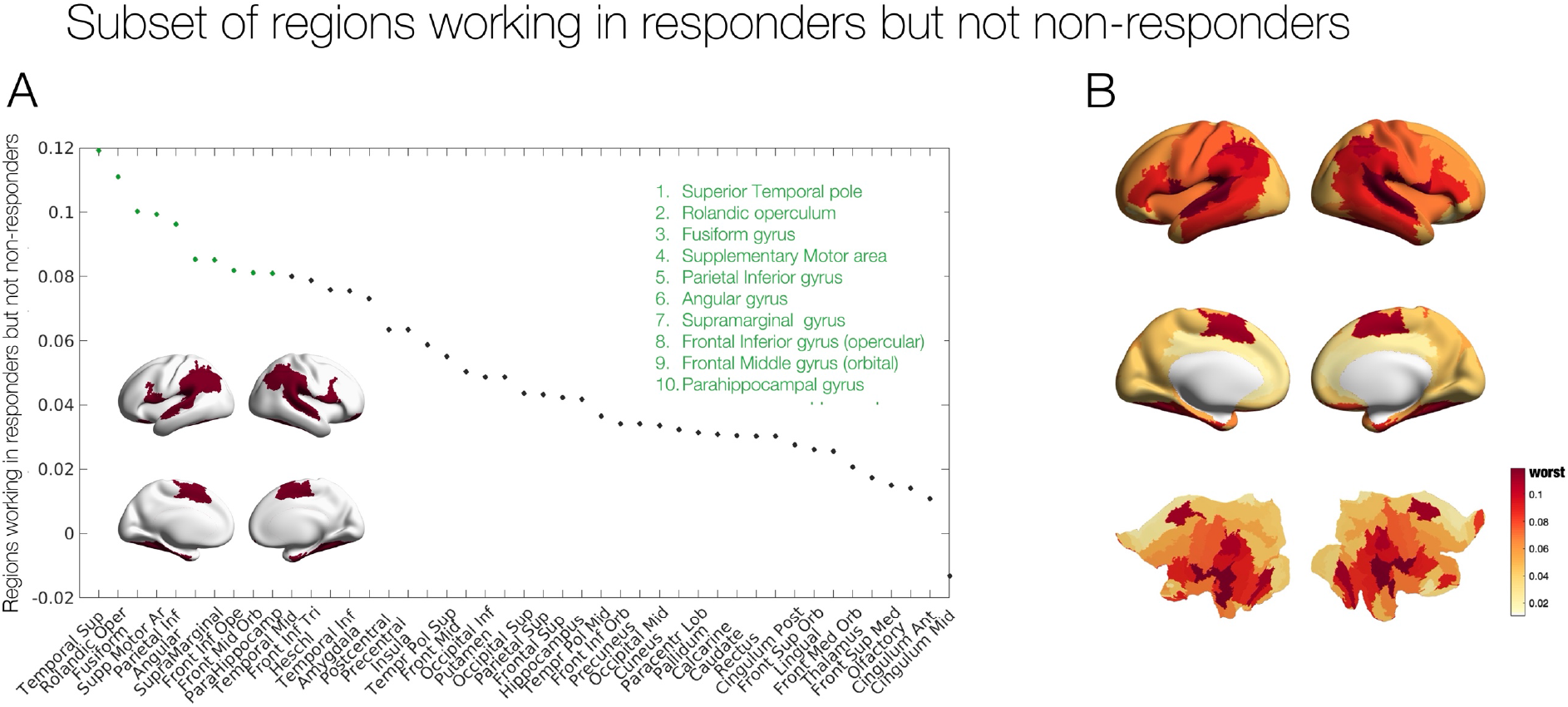
Subset of regions working in responders but not in non-responders. **A)** Rank ordered absolute difference of KL divergence between perturbations of the responder and non-responder models before treatment at a stimulation intensity of **a** = 0.07. Inset brain rendering of the ten brain regions with the highest difference: Temporal Superior pole, Rolandic operculum, Fusiform gyrus, Supplementary Motor Area, Parietal Inferior gyrus, Angular gyrus, Supramarginal gyrus, Frontal Inferior gyrus (opercular), Frontal Middle gyrus (orbital) and the Parahippocampal gyrus. **B)** Cortical rendering and flat maps showing the distribution of all KL divergence differences.

Figure 4.A shows the rank ordered regional differences in KL divergence between perturbations of the responder and non-responder models before treatment at a stimulation intensity of *a* = 0.07. We highlighted the regions with the largest KL divergence working in responders but not non-responders to promote a transition to the healthy state. These regions are the Temporal Superior pole, Rolandic operculum, Fusiform gyrus, Supplementary Motor Area, Parietal Inferior gyrus, Angular gyrus, Supramarginal gyrus, Frontal Inferior gyrus (opercular), Frontal Middle gyrus (orbital) and the Parahippocampal gyrus. Figure 4, Right, shows the cortical rendering of these differences.

### Correlation with Serotonin Receptor Maps

Given the unique neuropharmacology of the psychedelic-induced state through serotonergic receptors, we assessed whether the regions working in responders but not non-responders overlapped with the 5-HT density maps derived from PET imaging data previously obtained by an independent research group (Beliveau *et al*., 2017). Figure 5 A shows correlations between the *5-HT*_*2a*_ and *5-HT*_*1a*_ receptor density maps and the KL divergence differences for the two groups at optimal *a* = 0.07 (Spearman *ρ* = 0.227, p = 0.032 and Spearman *ρ* = 0.284, p = 0.007 respectively). Figure 5 B, shows non-significant correlations to other 5-HT components – namely the *5-HT*_*2b*_ (Spearman *ρ* = 0.064, p = 0.055) and *5-HT*_*4*_ receptors (Spearman *ρ* = 0.055, p = 0.607) plus the 5-HT transporter (*5-HTT*) (Spearman *ρ* = - 0.172, p = 0.106).

**Figure 5.**
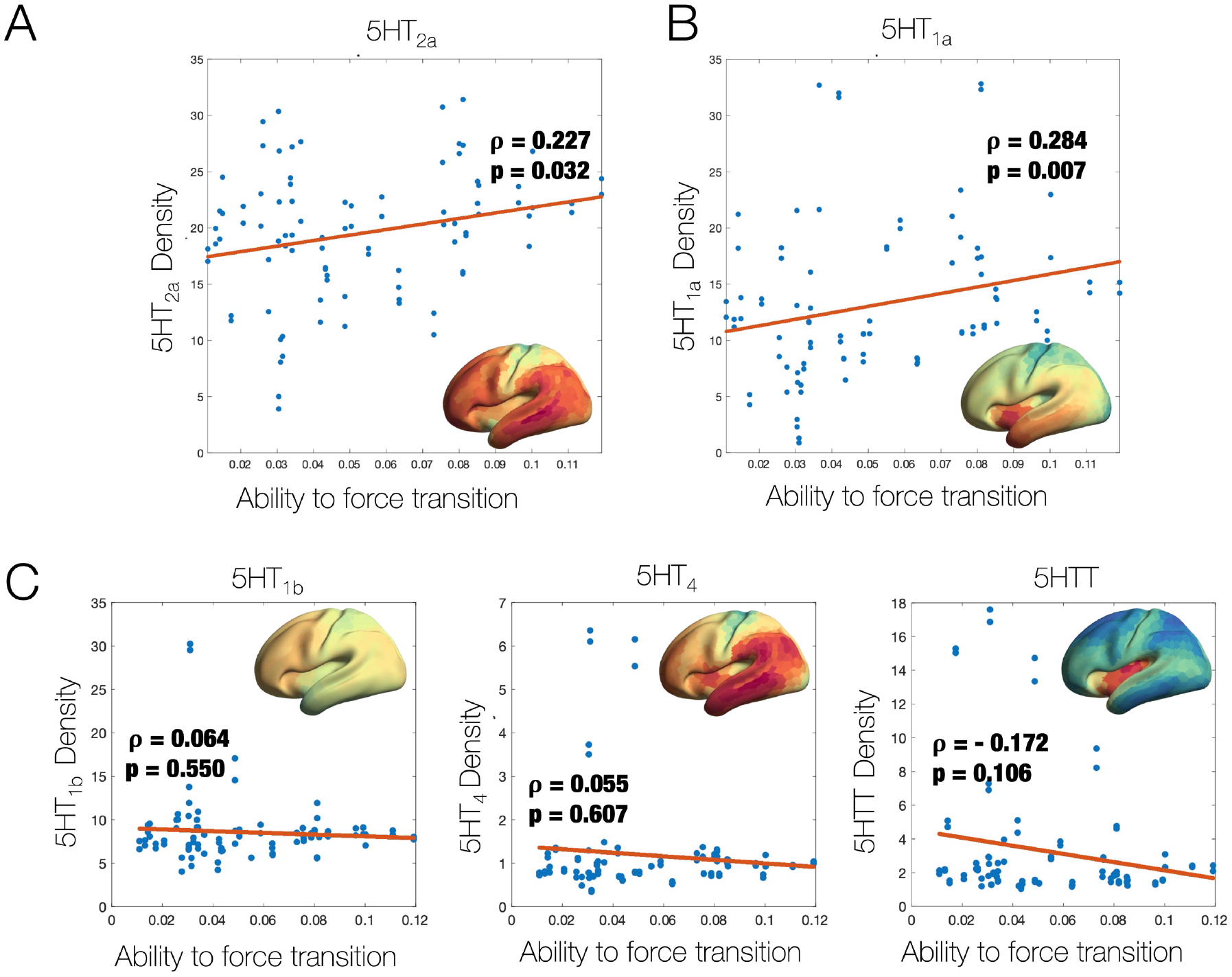
Ability to promote a transition relates to density of specific serotonin receptors: For each pair of homological brain regions, the ability to promote a transition is plotted against the receptor map densities of: **A)** 5-HT_2a_ (Spearman correlation ρ= 0.227, p = 0.032). **B)** 5HT_1a_ (ρ= 0.284, p = 0.007), and **C)** for other 5-HT receptors with non-significant results: 5-HT_2b_ (ρ= 0.064, p = 0.055), 5-HT_4_ (ρ= 0.055, p = 0.607) and the 5-HTT (ρ= -0.172, p = 0.106).

## Discussion

In this work, we employed a whole-brain modelling approach to evaluate potential brain-change causes of response to psilocybin therapy for treatment-resistant depression. Using a novel combination of empirical data and in silico modelling, systematic perturbations to brain regions modelled in silico, revealed a subset of regions implicated in transition away from ‘depressed brain’ pathology and towards the ‘healthy brain’ configurations of treatment responders. Notably, these regions matched those with the highest density of *5HT*_*2a*_ and *5HT*_*1a*_ neuroreceptors. This relationship is plausible given that psilocin (psilocybin’s active metabolite) is known to have an appreciable-to-high affinity for the 5-HT1A and 2A receptors, respectively, where it acts as an agonist; in the case of the 5-HT2AR, potentially stimulating plasticity-related signaling cascades relevant to an antidepressant action (Desouza *et al*., 2021; Liu *et al*., 2022).

A summary of complex spatio-temporal dynamics, in terms of brain substates and their transitions, has drawn a lot of attention in systems neuroscience due to its utility to evaluate the impact of pharmacological and electromagnetic interventions for treating brain and behavioural disorders. Brain substates have been characterised in different ways; by minimal energy (Gu *et al*., 2018) as attractor landscapes (Deco and Jirsa, 2012; Vohryzek *et al*., 2020), and more heuristically, through sliding-window analysis and unsupervised clustering (Hutchison *et al*., 2013; Allen *et al*., 2014). However, it has been challenging to find a model that sufficiently simple and yet accurate to account for temporally and spatially complex and non-stationary datasets. Here, PMS are built on a description of the data in terms of a probabilistic “cloud” in substate space and as such can be extended to different neuroimaging modalities with higher temporal resolution, such as EEG and MEG, or potentially to more fine-grained spatial resolutions (Deco *et al*., 2019; Kringelbach *et al*., 2020).

Cutting-edge non-invasive brain stimulation techniques such as Transcranial Magnetic Stimulation (TMS) and Direct Electrical Stimulation (DES), and new neuropsychopharmacological drugs for treatment of psychiatric disorders have heralded a new era of localized brain perturbations as medical interventions. For example, TMS has been considered for treatment of many psychiatric disorders such as depression, schizophrenia and addiction (Ridding and Rothwell, 2007), and classic psychedelic (drug) therapy, which in part, targets a specific neuroreceptor (i.e., principally the 5-HT2A receptor) is showing efficacy in the treatment of a broad range of conditions such as depressive, anxiety and addiction disorders (Carhart-Harris and Goodwin, 2017). However, it seems highly likely that the mechanistic action of these interventions lies - potentially well downstream of their initial action, and this action may not be straightforward (Turkheimer *et al*., 2021). For example, how Direct Electrical Stimulation (DES) induced signal propagates within neuronal microcircuits remains unclear and often paradoxical (Logothetis *et al*., 2010), and motivates theoretical neuroscience studies and *in-silico* perturbation protocols where system-wide changes to localised external stimulation can be explored.

Beyond *in-silico* perturbations, the exhaustive stimulation protocol can also be used as a dynamic sensitivity analysis tool from the complex systems perspective. Traditionally, statistical differences in measures summarising spatio-temporal dynamics are obtained using signal detection theory. Such approaches can be enhanced by considering whole-brain models and their structural differences between conditions, for example as described by the global coupling (*G*) parameter. Moreover, rather than describing and assessing expressions of spatio-temporal dynamics, an exhaustive protocol allows a shift of focus onto transitions to a target state and this can be used to identify differences between groups, such as treatment responders versus non-responders, as we have done here.

Forcing transitions in large-scale brain networks has also been investigated through the prism of control network theory. In such scenarios, control strategies are deployed to navigate complex systems from a source (initial) state to a target (final) state (Srivastava *et al*., 2020). This approach has obtained a lot of attention due to its wide-ranging engineering applicability in technological, social and cyberphysical systems across various experimental scenarios (Gu *et al*., 2015; Lynn *et al*., 2020). However, the conceptual understanding of controlling neuronal signals from source to target might be problematic as the brain operates in self-sustained and non-equilibrium state, and the notion of well-defined pathway between them might be ill-posed (Tognoli and Kelso, 2013). On the contrary, the approach considered in this work describes spatio-temporal dynamics in terms of Probabilistic Metastable Substates and, through systematic perturbation, rebalances the spatio-temporal dynamics between two PMS spaces. Through this approach the brain is rebalanced to its healthy working point, without specific instructions of what the relevant pathway might be.

To obtain a PMS approximation of the brain substate of interest, several methodological choices are made which inevitably introduce several caveats. Firstly, a regional parcellation must be chosen, which might introduce artificial spatial boundaries especially when dealing with dynamics. Secondly, the choice of clustering algorithm defines the type of substates that can be obtained. Here, we use the unsupervised learning algorithm *k-means* clustering which has been shown to adequately represent functionally meaningful brain substates (Vohryzek *et al*., 2020)-However, alternative algorithms could be used for this purpose (e.g. k-medoids). Related to the experimental data, the design is an uncontrolled open-label feasibility pilot study, and as such has no placebo group and suffers from small sample size. Hence, future replication studies are warranted to ensure robustness of the findings. Moreover, the healthy state is defined here in terms of the 1-day post-treatment scan but the responders/non-responders’ assessment is done 5 weeks after. Lastly, the whole-brain models constructed are based on group approximations of the functional brain information and structural connectivity group template. For clinical relevance, further research will be needed to create individual-based whole-brain models that might allow for future in-silico assisted personalised psychiatry (Deco and Kringelbach, 2014).

## Material and methods

### Experimental Data

#### Functional MRI

We carried out the analysis on previously published dataset of patients with treatment-resistant depression undergoing treatment with psilocybin at Imperial College London (Carhart-Harris *et al*., 2016). In brief, we investigated 15 patients (without excessive movement and other artefacts from the original 19 patients) who were diagnosed with treatment resistant major depression. The MRI scanning sessions were completed pre-treatment with psilocybin and one-day post-treatment with the treatment consisting of two oral doses of psilocybin (10mg and 25mg, 7 days apart). The patients were split into responders and non-responders to the treatment based on the Quick Inventory Symptomatology (QIDS) at 5-weeks post-treatment with 6 out of the 15 patients meeting criteria for response (Carhart-Harris *et al*., 2017).

#### Structural Connectivity

In this study, white-matter (structural) connectivity of 90 AAL brain areas from a previously obtained dataset was used for the whole-brain network model. In brief, the group consisted of 16 healthy young adults (5 females, mean SD age: 24.7 ± 2.54). Diffusion Tensor Imaging (DTI) was applied following the methodology described in (Cabral *et al*., 2012). Undirected structural connectivity *C*_*np*_ was obtained were ***n*** and ***p*** are brain areas and the connectivity weights are defined as the proportion of sampled fibers in all voxels in region ***n*** that reach any voxel in region ***p***. Finally, the individually structural connectomes were averaged across the 16 subjects to obtain a group-based template.

#### Probabilistic Metastable Substates

Firstly, we calculated the instantaneous phased relationship between individual brain regions by expressing the demeaned regional fMRI signal *x*(*t*) as an analytical signal i.e. in terms of its time-varying phase *θ*(*t*) and amplitude *A*(*t*) as *x*(*t*) = *A*(*t*) ∗ cos (*θ*(*t*)) (Glerean *et al*., 2012). We excluded the first and last three timepoints to account for the boundary artefacts introduced by the Hilbert transform. Hence for every time point *t* and pair of brain regions *n* and *m*, we obtain the phase coherence matrix dPC as follows:

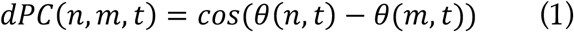

By decomposing the signal in this way, we can look at when the brain regions *n* and *m* are aligned with similar angles, *cos* (0) = 1, orthogonal to each other *cos* (*π/2*) = 1 and anti-aligned *cos* (*π*) = *−*1. As the phase coherence is a measure of undirected connectivity, the phase coherence matrix *dPC* is symmetric and all the meaningful information is captured in the upper-triangular matrix.

For further analysis, we used only the *1xN* leading eigenvector *V*_1_(*t*) of the dPC matrix as described in the Leading Eigenvector Dynamics Analysis (Cabral *et al*., 2017). In detail, at every timepoint *t* of the *dPC*(*t*), we performed the eigendecomposition taking the first (most dominant) eigenvector to describe the *dPC*(*t*) pattern. The *dPC*(*t*) is decomposed as *dPC*(*t*) = *V*(*t*)*D*(*t*)*V*^−1^(*t*) where D is the diagonal matrix carrying the real-valued eigenvalues and *V*_1_(*t*) and *V*^−1^_1_(*t*) are the left and right corresponding orthogonal eigenvectors respectively. The dominant connectivity pattern can be simply reconstructed by the following matrix multiplication *V*(*t*)*V*^−1^(*t*).

To look for and describe the discrete phase-locking states, we clustered all the leading eigenvectors obtained from all the fMRI scans obtained from both responders and non-responders. We used the unsupervised k-means algorithm, of varying cluster number *k* from 2 to 10 clusters, to iteratively converge to a predefined number of clusters with 20 random cluster initialisations to ensure stability in the clustering. Again, by computing the matrix multiplication of the *1xN* cluster centroids *V*_*cα*_ as 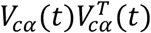 we obtain the dominant connectivity pattern of each cluster. In the current analysis, we considered the cluster solution *k* = 3 as an optimal choice between the quality measures - Dunns, Davies-Bouldin and Silhouette Score, Davies (SI Figure 2), and the maximising of the statistical significance between patient groups (p-values).

After calculating the phase-locking states, we defined the probability of occurrence of the individual substates by simply dividing their occurrence in each recording session by the total number of time points recorded (same for all recordings).

#### Whole-brain Computational Model

In order to simulate the ultra-slow fluctuations in fMRI signal detected during rest, we used the Landau-Stuart oscillator canonical model, describing the transition from a noisy to an oscillatory dynamics (Kuznetsov, 1996). The so-called supercritical Hopf-bifurcation model was used locally at every brain region (node) to emulate the local dynamics (Deco, Kringelbach, *et al*., 2017; Deco *et al*., 2019). To achieve a whole-brain level description, the individual Hopf models were coupled in a structural connectivity (SC) network, describing the large-scale white-matter map of the human brain (Hagmann *et al*., 2008; Deco, Kringelbach, *et al*., 2017). The emerging and complex interactions in the whole-brain network of coupled Hopf models have been shown to describe many aspects known from experimental recordings in MEG (Deco, Cabral, *et al*., 2017) and fMRI (Kringelbach *et al*., 2015; Deco and Kringelbach, 2016; Deco, Kringelbach, *et al*., 2017; Deco, Cabral, *et al*., 2018; Deco *et al*., 2019).

Formally, the normal form of the supercritical Hopf-bifurcation model for a single uncoupled region of interest (*n*) in Cartesian coordinates is described by the following set of coupled equations:

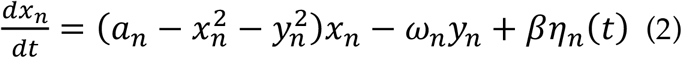

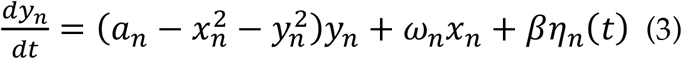

with *βη*_*n*_(*t*) being the gaussian noise with standard deviation of *β* = 0.02. The bifurcation parameter ***α*** positions the system at the supercritical bifurcation point when ***α*** = 0, noise activity governed by *βη*_*n*_(*t*) in regime when ***α*** < 0, and stable limit cycle with oscillatory behaviour of frequency defined by *f*_*n*_ = *ω*_*n*_/2*π* when ***α*** > 0. The values of the intrinsic frequency *ω* were calculated from the experimental fMRI signals in the 0.04 *−* 0.07 *Hz* band by taking the peak frequency of the gaussian-smoothed power spectrum of each brain area.

To describe the coupled whole-brain computational model, we introduced the coupling term (modelled as the common difference coupling i.e., describing the linear term of a general coupling function) between the individual nodes weighted by the corresponding values of the SC matrix. To be noted, we do not consider the next nonlinear coupling term following Taylor expansion of the full coupling, in case the linear coupling is non-existent (Kuramoto, 1984; Pikovsky *et al*., 2002). The equations 2 and 3 can be hence expanded as follows:

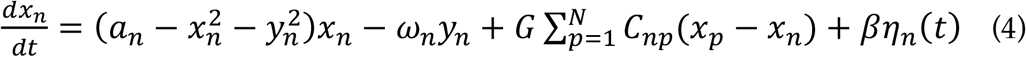

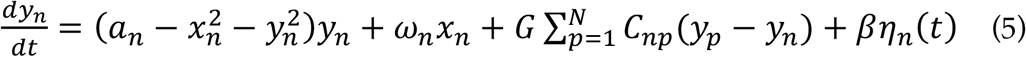

where *C*_*np*_ is the SC weight between node *n* and *p*, and *G* is the global coupling weight with equal contribution between all the nodal pairs. The SC matrix was rescaled to have the mean value < *C* > = 0.2 in order to be consistent with previous literature’s range of parameters (Deco, Kringelbach, *et al*., 2017; Deco *et al*., 2019). The simulated signal is described by the *x*_*n*_ equation for every node *n*. The variables *G* and *a* are the control parameters used for the model fitting to the experimental data and the stimulation protocol respectively (Deco, Kringelbach, *et al*., 2017; Deco *et al*., 2019).

#### Objective Function

In order to validate the simulated signal different realisations of the experimental data can be used (Cabral, Kringelbach and Deco, 2017). The most standard approach is comparison of the simulated data with grand-averaged static functional connectivity as computed by the Pearson correlation (Honey *et al*., 2009; Deco and Jirsa, 2012) or metastability defined as the standard deviation of the Kuramoto Order Parameter (SI Figure 4 - Metastability). To account for the temporally varying nature of the BOLD signal, recent literature has focused on the comparison between the simulated and empirical FCD spectrums (quantified by Kolmogorov-Smirnov distance) i.e. the distributions of the cosine distance between the consecutive timepoints as described by the leading eigenvector (Deco, Kringelbach, *et al*., 2017; Deco, Cruzat, *et al*., 2018) (SI Figure 4 – Functional Connectivity Dynamics). As alluded to in the previous section, the fMRI signals organise into spatially meaningful phase-locking states. Here, we compare the simulated data to the probabilities of occurrence of the phase-locking states found in the experimental recordings (Deco *et al*., 2019). We used the symmetrised Kullback-Leibler Divergence (KL divergence) of the simulated and empirical probabilities of occurrence as follows:

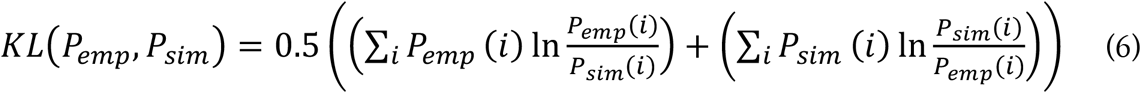

with *P*_*emp*_ and *P*_*sim*_ being the empirical and simulated probabilities of occurrence of the same phase-locking states respectively.

## Acknowledgements

M.L.K. is supported by the European Research Council Consolidator Grant: CAREGIVING (615539), Pettit Foundation, Carlsberg Foundation and Center for Music in the Brain, funded by the Danish National Research Foundation (DNRF117). Joana Cabral is supported by Portuguese Foundation for Science and Technology CEECIND/03325/2017, UIDB/50026/2020 and UIDP/50026/2020, Portugal. Gustavo Deco is supported by the Spanish Research Project PSI2016-75688-P (Agencia Estatal de Investigación/Fondo Europeo de Desarrollo Regional, European Union); by the European Union’s Horizon 2020 Research and Innovation Programme under Grant Agreements 720270 (Human Brain Project [HBP] SGA1) and 785907 (HBP SGA2); and by the Catalan Agency for Management of University and Research Grants Programme 2017 SGR 1545.

