## Supplementary Information for "Brain dynamics predictive of response to psilocybin for treatment-resistant depression"

### When psychedelics work for depression: Insights from brain dynamics in responders and non-responders

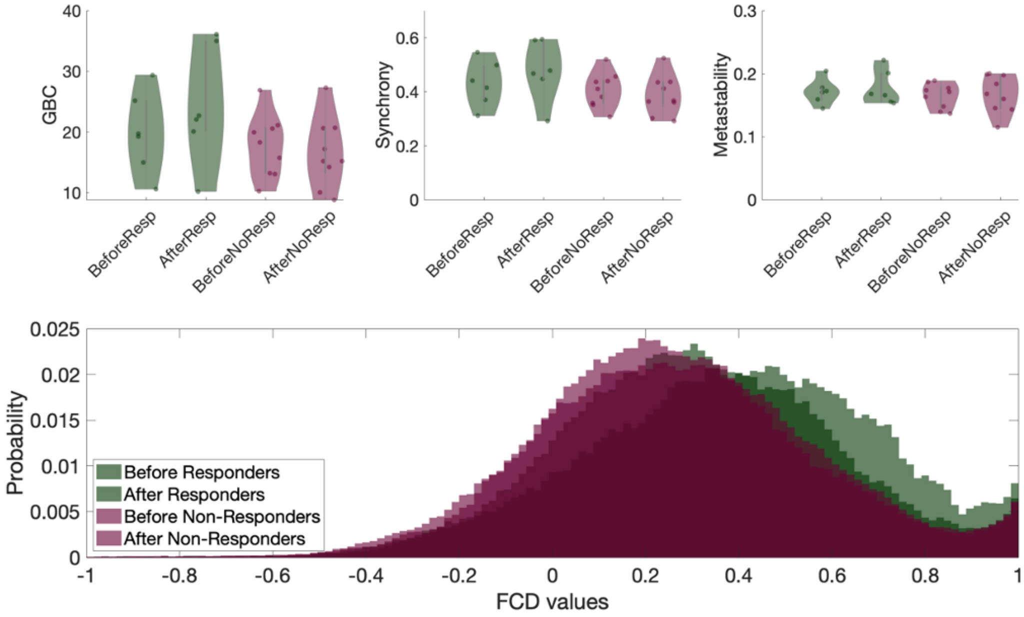


***SI Figure 1. Empirical Results - GBC, Synchrony, Metastability, FCD.*** *Results of other measures between responders and non-responders to the psilocybin treatment. Here, non-significant differences reported for Global Brain Connectivity (GBC), Synchrony and Metastability. Furthermore, the mean histogram values of the FCD spectrum were found to be significant between pre- and post-treatment responders (p = 0.0163, signed rank-sum permutation test), and pre- and post-treatment non-responders with post-treatment responders respectively (p = 0.0183 and p = 0.0273, rank-sum permutation test), demonstrating the importance of taking into the consideration temporal fluctuations.*

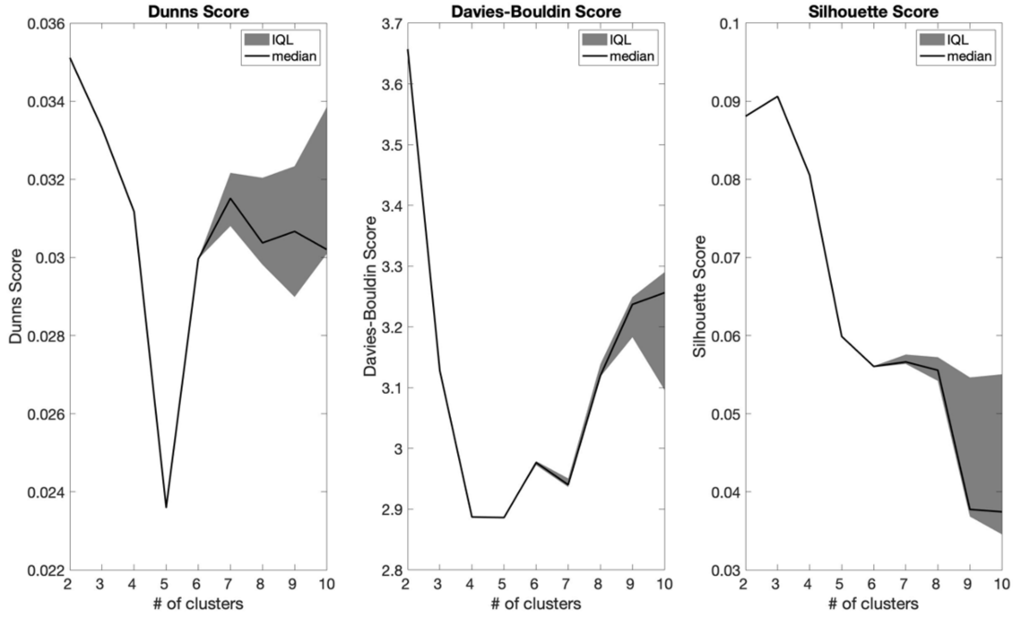


***SI Figure 2. Clustering Quality Measures.*** *Three quality measures for the clustering solutions across runs of varying number of clusters. Here, Dunn's score, Davies-Bouldin and Silhouette scores are reported.*

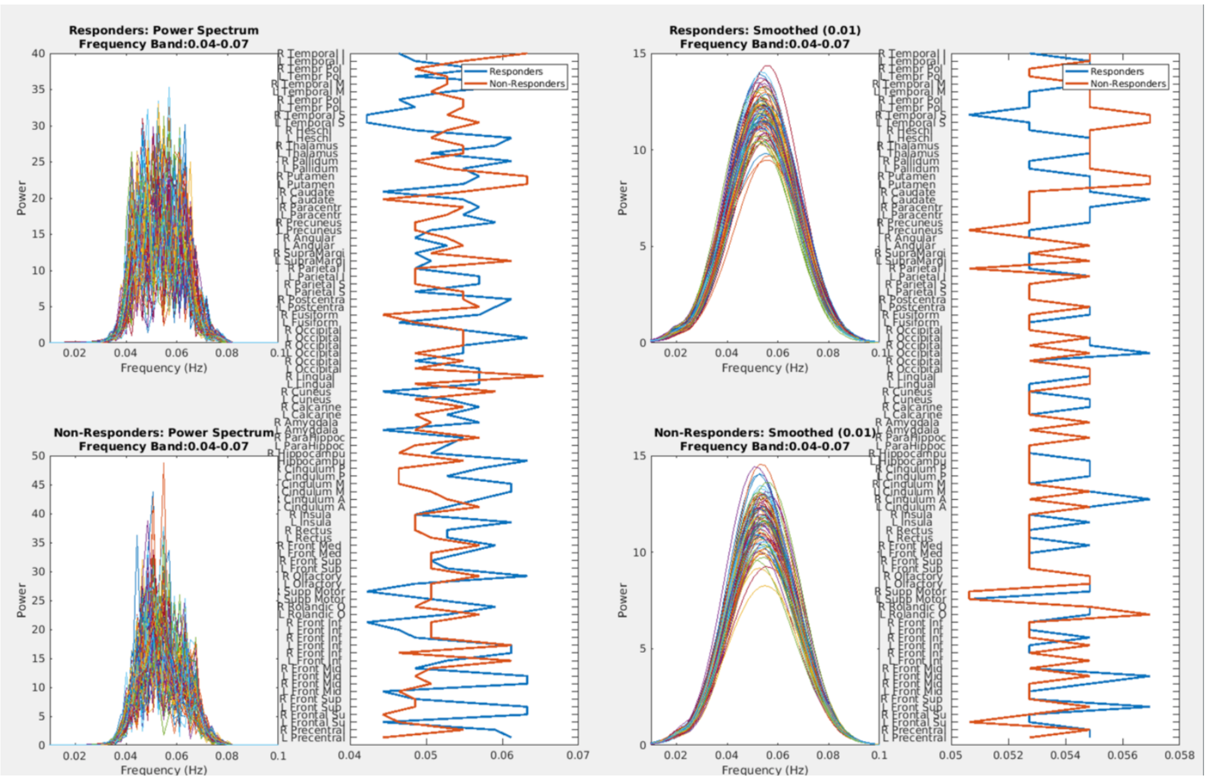


***SI Figure 3. Frequency Analysis of the experimental dataset.*** *Power Spectrum Distribution for responders and non-responders for unsmoothed (Left) and unsmoothed power spectrum. The responders and non-responders Hopf models are constructed to account for regional frequency heterogeneity of the smoothed Power Spectrum Distributions.*

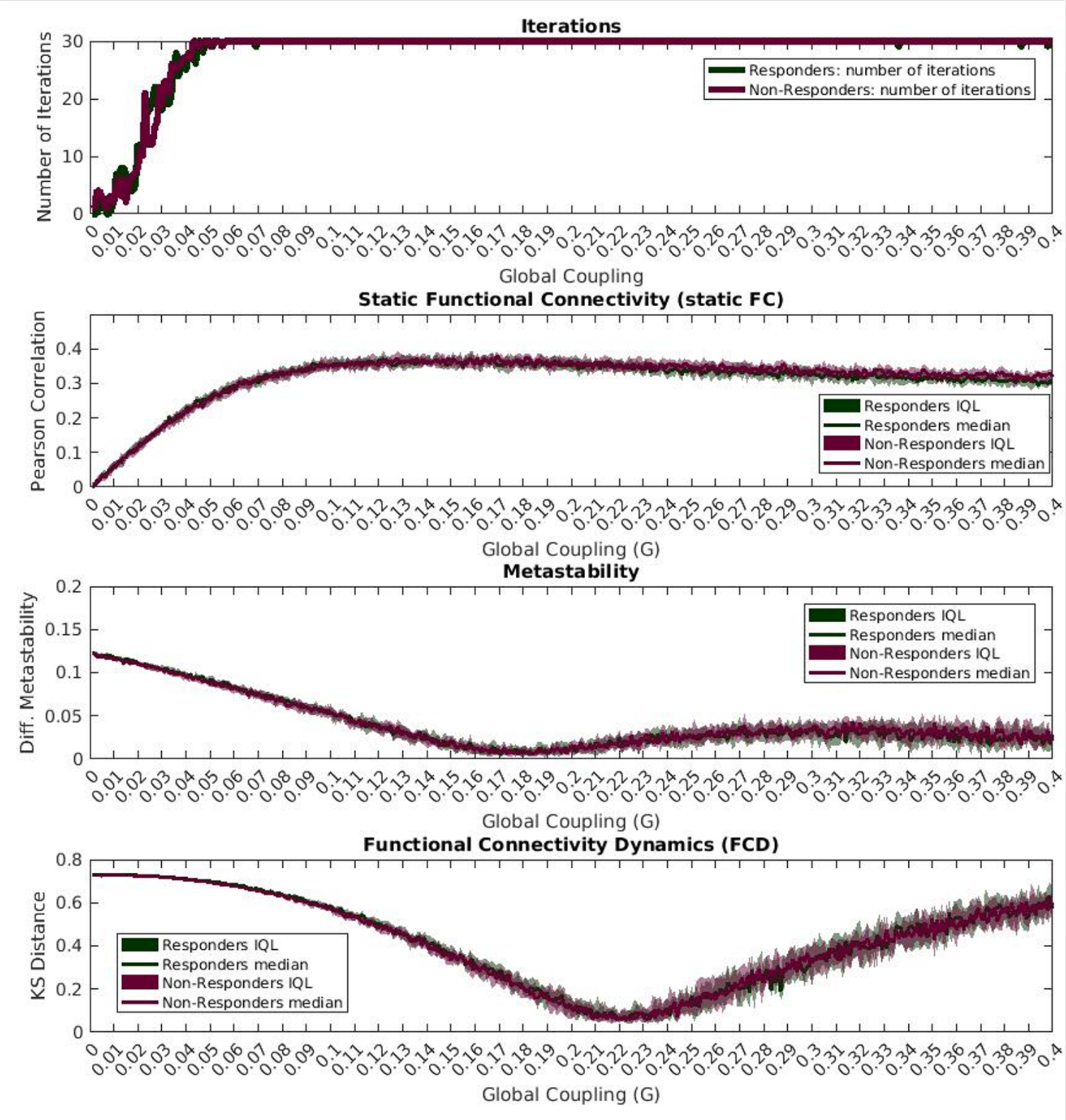


***SI Figure 4. Model Validation - GBC, Synchrony, Metastability, FC.*** *The model validation was run over 30 iterations. Results for other objective functions: static FC (Pearson Correlation), Metastability (Difference in metastability) and Functional Connectivity Dynamics (KS-distance).*
